# Evolutionary and Structural Bioinformatics Reveal GPR89 as a Conserved Solute Carrier Transporter

**DOI:** 10.1101/2025.08.04.668498

**Authors:** Camila A. Quercia-Raty, Luka Robeson, Sebastian E. Brauchi

## Abstract

Also called the Golgi pH regulator (GPHR), GPR89 is an orphan membrane protein found in nearly all eukaryotic lineages. Despite its broad phylogenetic distribution, the evolutionary history, structural diversity, and function of GPR89 remain poorly understood. In this study, we present a comprehensive bioinformatic analysis of GPR89 in Eukarya by integrating phylogenetic reconstruction, genomic synteny, sequence conservation, and structural modeling. While GPR89 is typically encoded as a single-copy gene, we identified lineage-specific duplications in both vascular plants and vertebrates. In contrast to the large sequence conservation, large differences can be observed in whether plants and animals preserve the gene structure flanking GPR89. Structural phylogenetic clustering places GPR89 within the solute carrier (SLC) superfamily, supporting previous reports claiming transporter-like activity rather than GPCR signaling. Predicted structures reveal a unique intracellular hairpin loop and a conserved transmembrane core compatible with transport activity. Our findings provide a unifying framework for interpreting the divergent functional roles reported for GPR89 and establish it as a structurally conserved, evolutionarily stable member of the SLC superfamily.

## INTRODUCTION

GPR89 is a highly conserved orphan membrane protein found across unicellular eukaryotes, land plants, and animals (Maeda et al., 2008; Charroux and Royet, 2013; Deckstein et al., 2015; Dong et al., 2019; Rojas et al., 2019). Deficient GPR89 function in mammals has been associated with neurodegeneration (Sou et al., 2019), disrupted brain cholesterol synthesis (Sou et al., 2022), and compromised epidermal barrier function (Tarutani et al., 2012). Originally annotated as a G protein-coupled receptor (GPCR), GPR89 displays nine predicted transmembrane domains, differing from the canonical seven-transmembrane GPCR structure. Although interactions with the G protein alpha subunit have been reported in plants such as rice, maize, and *Arabidopsis thaliana* (Pandey et al., 2009; Ma et al., 2015; Dong et al., 2019; Zhang et al., 2024), GPR89’s classification as a functional GPCR remains unconfirmed due to the absence of a defined activation mechanism or details regarding a preferent signaling pathway (Jaffé et al., 2012).

Interestingly, GPR89 has been linked to a variety of molecular functions that might seem incompatible at first. In vascular plants, its localization and proposed roles vary widely; it has been described as a calcium-permeable channel activated by cold (Ma et al., 2015), an abscisic acid receptor (Pandey et al., 2009; Zhang et al., 2024), and an ion channel present in the ER, Golgi, and the plasma membrane (Jaffé et al., 2012).

In animals, GPR89 has been referred to as the Golgi pH regulator (GPHR) and is predominantly located in the Golgi apparatus and endoplasmic reticulum (Maeda et al., 2008; Charroux and Royet, 2013; Sou et al., 2019). A particularly intriguing functional hypothesis suggests that mammalian GPR89 forms a large-conductance, non-rectifying chloride channel (∼300 pS; P_Cl-_/P_K+_ ≈ 2.5) in the Golgi membrane, potentially corresponding to the GOLAC-2 current (Maeda et al., 2008; Thompson et al., 2002). This chloride conductance may provide the counterion flux necessary to support proton accumulation and maintain luminal pH gradient in Golgi. Importantly, Golgi pH regulation is associated with several pathologies, including congenital glycosylation disorders, cancer, viral susceptibility, and intellectual disabilities (Rivinoja et al., 2012; Khosrowabadi and Kellokumpu, 2020).

Lastly, in the unicellular eukaryote *Trypanosoma brucei*, GPR89 is found at the plasma membrane, where it likely functions as an oligopeptide transporter (Rojas et al., 2019). These divergent functions and localizations suggest that the different orthologs of GPR89 may have evolved distinct physiological roles across taxa, while maintaining a conserved core structure.

In this study, we investigated the evolutionary and structural landscape of GPR89 to better understand its conserved roles across eukaryotic lineages. We show that it is retained as a single-copy gene in most taxa, with independent duplication events observed in both mammals and vascular plants. Lineage-specific retro-pseudogenization is found predominantly in Anthropoid primates. Phylogenetic and structural analyses reveal that GPR89 shares sequence and structural homology with solute carrier transporters (SLC), clustering near the SLC39 and SLC42 families associated with zinc and ammonium transport, respectively. Mapping sequence conservation and evolutionary covariance in AlphaFold2-predicted structures highlights a conserved hydrophobic core centered on transmembrane segment 5 (TM5), forming part of a lateral groove that could be potentially involved in substrate binding. Surrounding this core are polymorphic intracellular and extracellular loops, including a prominent cytosolic coiled-coil hairpin (HP) structure. The loop at the tip of the hairpin (HPloop) shows marked sequence variability, especially among unicellular taxa, and may represent a modular domain contributing to the observed functional divergence of GPR89 across eukaryotic lineages. Overall, GPR89 represents a structurally conserved SLC-type transporter that likely acquired lineage-specific functional specializations, while retaining a core architecture associated with membrane permeability and organellar homeostasis.

## RESULTS

### Repertoire of *GPR89* across species

To understand the process of diversification of *GPR89*, we first performed a Maximum likelihood (ML) phylogenetic analysis using CDS from representative members of diverse taxonomic groups. Following manual annotation and calculation of hydrophobicity profiles, we recovered a total of 130 sequences, annotated as *bona fide GPR89* orthologs in model species (Supplementary Table 1). The collected orthologs had an average length of 1,368 nucleotides and the final alignment spanned 3,913 nucleotides, displaying 248 conserved sites.

The Maximum Likelihood tree obtained shows the presence of the *GPR89* gene among Metazoa, Archaeplastida, and Heterotrophic protists corresponding to Amorphea (Holozoa, Ascomycota, and Amoebozoa), Excavata, and SAR groups (Rhizaria and Alveolata) (Figure 1). The gene tree is in agreement with previous phylogenetic hypotheses obtained by unrooted Neighbor-Joining trees (Jaffé et al., 2012; Deckstein et al., 2015; Mojib and Kubanek, 2020). Our thorough phylogenetic reconstruction, recovers the monophyly among Metazoa, within Vertebrata, and its natural relations as monophyletic clades (Mammalia, Sauropsida, Anura, and Actinopterygii). Furthermore, the tree recovers Chlorophyta species as a sister group of Embryophyta (Figure 1). Other relationships—such as those established among heterotrophic protists—appear as paraphyletic clades, including organisms from SAR and Opisthokonta (Figure 1). Therefore, our analysis is consistent with both previous observations and the updated hypotheses for the speciation of Eukaryotic organisms (Burki et al., 2020).

**Figure 1:**
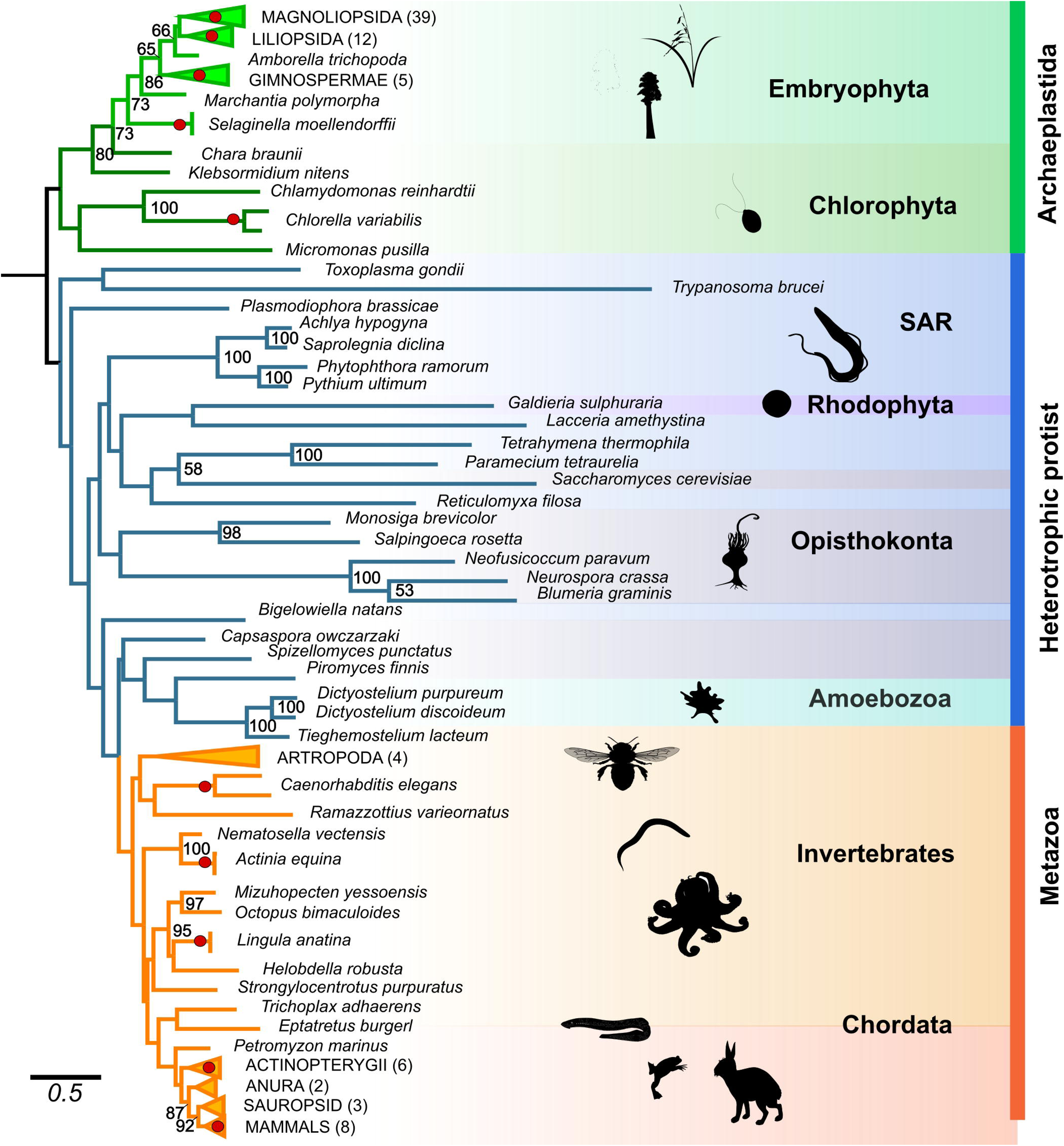
Duplication events of GPR89 across Eukarya. Best Maximum Likelihood phylogenetic tree displaying large groups in color code, Metazoa in orange, Archaeplastida in green, and heterotrophic protist in blue. Numbers under the nodes represent bootstrap support values and red circles denote GPR89 intraspecific functional gene duplication. Numbers in parenthesis indicate the sequence number for each collapsed clade. Bar indicates nucleotide substitutions per site. The tree was rooted at the midpoint.

The branch lengths within major clades are relatively uniform, which supports a high degree of sequence conservation among closely related taxa (Figure 1). An exception to this pattern is observed in heterotrophic protists, where significantly longer branch lengths indicate diversification. Among them, *Trypanosoma brucei emerges* as the most divergent, contrasting to the higher conservation observed within Metazoa and Archaeplastida.

A functional paralog of *GPR89A*, *GPR89B*, was found present in both Metazoans and Archaeplastida, but absent in heterotrophic protists. This duplication appears in several species among Metazoa, including members in Mammalia, Actinopterygii, Nematoda, Cnidaria, and Braquiopoda. On the other hand, green algae (*Chlorella variabilis)* and many species of vascular plants such as Lycophyte, Gymnosperms and Angiosperms (both Eudicotyledons and Monocotyledons), present two or more functional copies of this gene (Figure 1, red dots; Supplementary Figure 1).

More duplication events of the functional GPR89 gene were found in Archaeplastidia than in Metazoa, which was expected due to the polyploidization events occurring commonly in plants (Soltis and Soltis, 2016; Liu et al., 2021; Levy and Feldman, 2022; Zhang et al., 2022). We infer that the observed gene duplications across taxa did not originate from a single ancestral duplication event, but rather resulted from multiple independent duplication events (Supplementary Figure 1).

### Evolutionary trajectory of GPR89 among mammals

In addition to functional duplication, we identified a reported human pseudogene annotated as GPR89P. This drives a broader search for GPR89A-related pseudogenes. Among eukaryotes, we found GPR89P restricted to mammals, particularly in most Anthropoid primates, and sporadically in Lagomorpha (rabbit), Scandentia (treeshrews), Rodentia (naked mole rat, degu, jerboa), Cetartiodactyla (blue whale, dolphin), and Marsupialia (wombat, koala). Thus, to explore the evolutionary trajectory of GPR89 in mammals, we conducted a phylogenetic analysis including annotated sequences for GPR89A, GPR89B, and GPR89P (Supplementary Table 2). The resulting gene tree confirmed one-to-one orthology of GPR89A across mammals (Supplementary Figure 1A), with expected relationships among major lineages. An exception was a paraphyletic Rodentia, likely due to long-branch attraction involving *Jaculus jaculus* and *Dipodomys ordii* (Felsenstein, 1978; Bergsten, 2005). Our topology recovered monophyletic clades for Primate, Dermoptera, Scandentia, Lagomorpha, Atlantogenata, Carnivora, Cetartiodactyla, Chiroptera, Marsupialia, and Monotremata. Notably, in *Homo sapiens* and *Vombatus ursinus*, the GPR89A and GPR89B paralogs showed strong support as sister genes (bootstrap 100), suggesting species-specific duplications rather than a shared ancestral event (Supplementary Figure 1A). In Anthropoids, functional GPR89 and GPR89P formed separate monophyletic groups. Most pseudogenes lack introns, contain multiple insertions, deletions, and premature STOP codons, all hallmarks of retrotransposition (Zhang, 2003; Ciomborowska-Basheer et al., 2021). Exceptions were *Nomascus leucogenys* and *Rhinopithecus roxellana*, whose pseudogenes encode full-length proteins with fewer exons than the original gene. Pseudogenes were consistently located on different chromosomes, further supporting a retrotransposition origin. This likely reflects independent pseudogenization events across mammalian lineages (Mighell et al., 2000; Zhang and Gerstein, 2004), and would be consistent with differential gene retention during GPR89 evolution (Shiu et al., 2006; Cañestro et al., 2013; Albalat and Cañestro, 2016).

### Patterns of shared synteny in *GPR89* orthologs

To better understand the evolutionary history of GPR89, especially in groups with known duplication events and similar evolutionary time frames, we inspected their genomic organization. For this purpose, we compared the synteny found from from *H. sapiens* to *Danio rerio* (429 MYS of divergence; Kumar et al., 2022) (Figure 2A) and that of *A. thaliana* to *Anthoceros agrestis* (451 MYA of divergence; Kumar et al., 2022) (Figure 2B).

**Figure 2:**
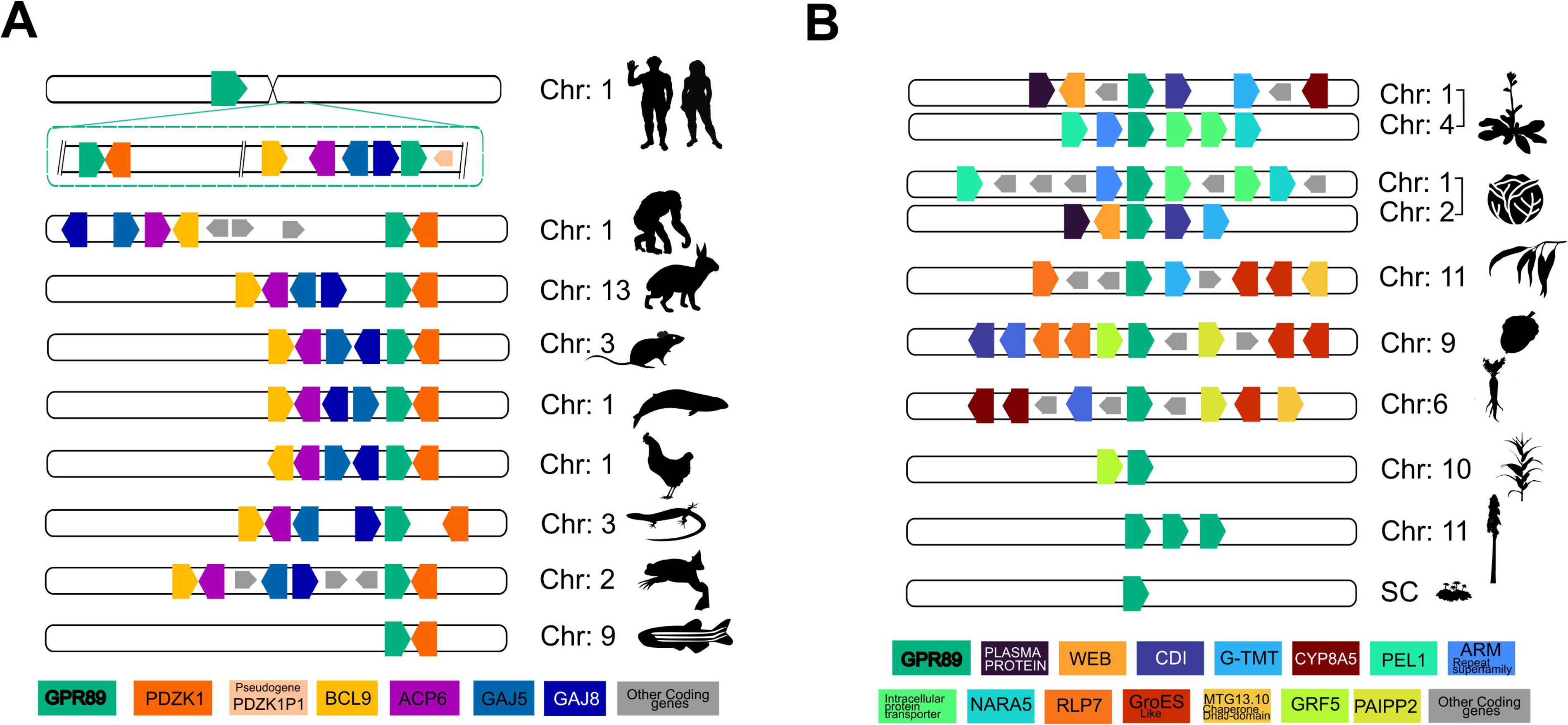
Synteny of the GPR89 locus in vertebrates and land plants. Conserved neighboring genes are shown as colored areas pointing at their transcriptional orientation. Corresponding chromosomes for each of the selected species is indicated. **(A)** Shows functional human GPR89 paralogs (GRP89A and GPR89B), both present in chromosome 1, together with GPR89 gene location among other metazoa species. **(B)** Displays *Arabidopsis thaliana* and *Brassica oleracea* paralogs on different chromosomes, together with other genes until Antocheros.

For the case of vertebrates, a comparison of the genomic regions displayed by the mammalian, sauropsid, and amphibian orthologs indicates that the single copy gene is flanked by five conserved genes: *GJA8*, *GJA5*, *APC6*, and *BLC9* upstream, and by *PDZK1* downstream. In fish, it is flanked downstream only by *PDZK1* (Figure 2A). The synteny observed in vertebrates is only partially conserved in humans. The two functional copies of GPR89 arose in human lineage from a unique duplication event that happened approximately 4.7 million years ago (Dennis et al., 2017). These copies are located in close proximity to each other on chromosome 1 (loci: 1:145,670,650-147,993,592) (Figure 2A). In the case of *GPR89A*, it is flanked only by *PDZK1* downstream. Meanwhile, *GPR89B* is flanked by *GJA8*, *GJA5*, *APC6*, and *BLC9* upstream, and a pseudogene of *PDZK1* is now present downstream (Figure 2A). In contrast to the functional genes, human *GPR89P* is present on chromosome 6 (loci: 6:27.737.000-27738.494) and, consistent with its nature as a retropseudogene (Zhang, 2003; Ciomborowska-Basheer et al., 2021), it is not flanked by any of the aforementioned genes. In other words, the origin of *GPR89B* and the appearance of *GPR89P,* in humans, represent independent events during the evolution of the gene. Based on our phylogenetic reconstruction and synteny analysis, it is evident that a second functional copy resulting from the duplication of GPR89 in Mammalia is only present in two species, whereas the retropseudogene (GPR89P) exhibits a higher frequency in the mammalian clade and, according to databases, encodes a putative protein in two primate species.

In contrast, Embryophyta displays a similar synteny only at family taxonomic level (Figure 2B). For instance, in Brassicales, *A. thaliana* GPR89A (known as AtGTG1) on chromosome 1 (loci: 1:24,139,851-21,146281) has similar gene composition than his orthologous in *B. oleracea* on chromosome C2 (loci: C2:14,525,378-14530,132). Similar observation occur with *A. thaliana* GPR89B (known as AtGTG2) on chromosome 4 (loci: 4:13,791,642-13,797,305) and *B. oleracea* GPR89B on chromosome C1 (loci:14,102,087-14,105,971). However, among different taxonomic orders, similarities can be observed in a limited number of instances. For example, *E. grandis* and *Q. lobata* (Myrtales order) shared GroES Like and WEB genes on the same chromosome where GPR89 is located, but in different positions and orientations. Similar observations were found between *Q. lobata*. *D. carota*, which shared, in different positions and direction, ARM Repeat superfamily, PAIPP2, and GroES-like genes. It is noteworthy that the three copies found in *Sequollodendron giganteum* and the two copies of *Solanum tuberosum,* are present in the same chromosome as a tandem duplication (Figure 2B and Supplementary Table 2).

Both synteny and phylogenetic analyses revealed that extra GPR89 copies appear on homologous or different chromosomes, consistent with polyploidy or retrotransposition respectively, regardless of taxonomic group (Panchy et al., 2016). Despite comparing similarly divergent taxa, synteny in chordates is remarkably conserved, unlike the extensive genomic rearrangements seen in plants, a difference that likely reflects the higher tolerance of plants to chromosomal reorganization.

### Amino acidic conservation patterns highlight critical regions

In order to extract meaningful information from the amino acid sequence, we obtained protein sequences by translating our annotated set of sequences (Supplementary table 1). The final set included representatives of Opisthokonta (Metazoa, Fungi, and Choanozoa), Chlorophyta (land plants and single-celled algae), as well as heterotrophic protists, such as members of the groups SAR, Rhodophyta, and Amoebozoa. We employed amino acid frequency histograms to detect conserved and variable regions within the protein sequence of human GPR89 (Figure 3). We observed that the regions encoding for TM segments previously suggested in the literature (Maeda et al., 2008; Pandey et al., 2009; Charroux and Royet, 2013; Dong et al., 2019; Zhang et al., 2024) are not only well defined but largely conserved among all taxa (Figure 3A). A thorough analysis of the amino acid distribution revealed 4 residues with conservation over 95%, 3 of which are present within the region corresponding to putative TM5 (Figure 3, A & B). Moreover, two regions of comparatively high variability exist within the linkers flanking the conserved TM5 (Figure 3C).

**Figure 3:**
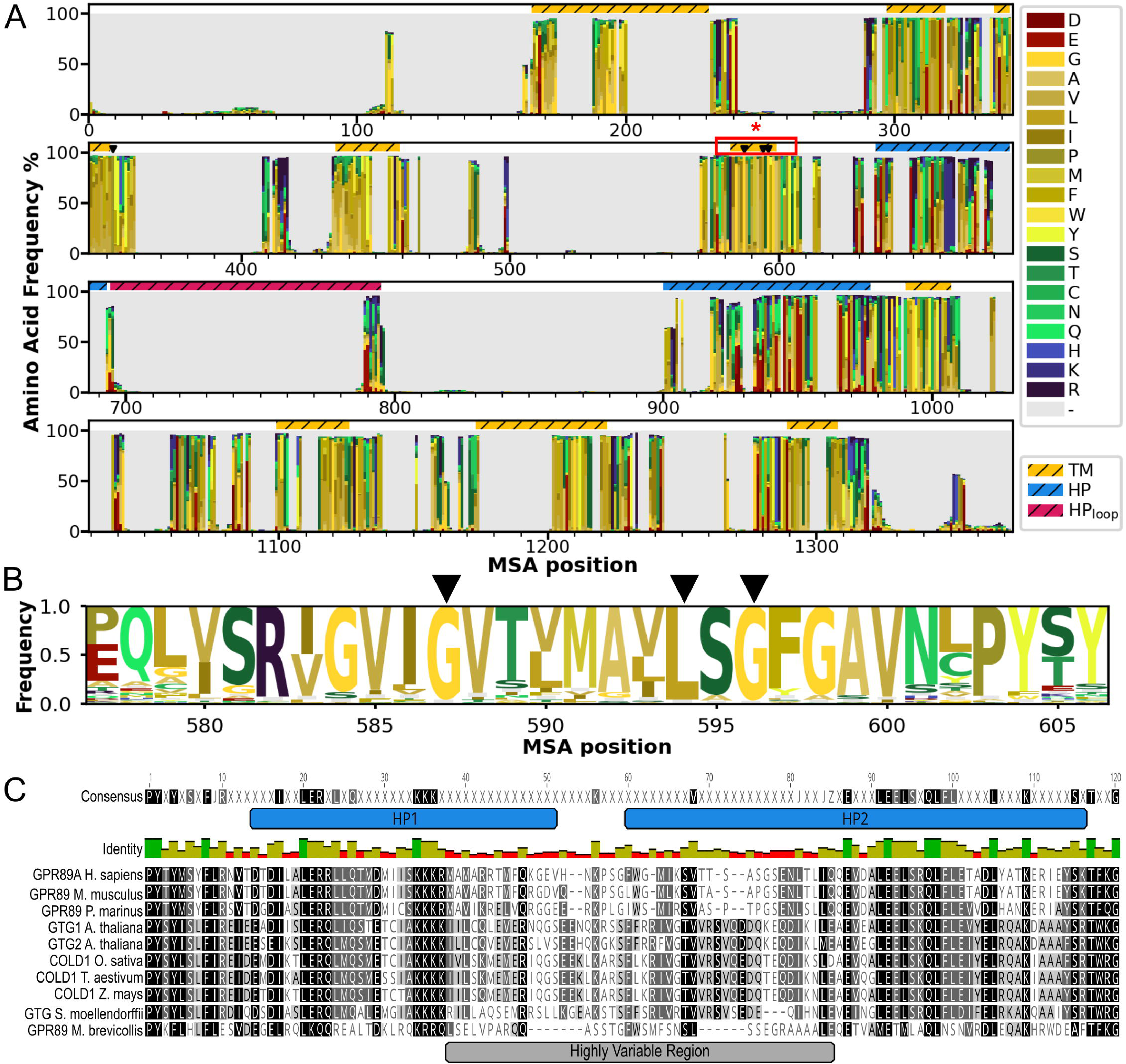
Multiple Sequence Alignment of the translated sequences of curated GPR89 orthologs found across Eukarya. **(A)** Per-residue frequency histogram for the different amino acids. Structural annotations from the human GPR89A protein, obtained from the AlphaFold DB and DeepTMHMM, are mapped to their corresponding positions within the histogram: transmembrane helices (TM1-9, gold/hatched), hairpin helices (HP1&2, blue/hatched), and hairpin loop (magenta/hatched). Black triangles above the bars mark the positions of highly conserved residues (>95%). A red rectangle delimits a cluster of highly conserved residues at TM5. **(B)** Amino acid frequency logo of the TM5 conserved cluster. **(C)** MSA of the hairpin region of selected GPR89 orthologs from animals, plants, and one choanoflagellate. Residue similarity is indicated at 100% (black), >80% (dark gray) & >60% (light gray). A consensus sequence of residues over 90% conserved is shown on top. A highly variable region is observed surrounding and encompassing the hairpin loop.

An important degree of polymorphism, including a variety of insertions/deletions and natural mutations, was found in the portion of the intracellular linker located between TM5 and TM6 (Figure 3, A & B). The most dramatic changes in the polymorphic region correspond to large insertions that are present in unicellular eukaryotes such as Amoeba, green algae, Ascomycota, and Excavata. Interestingly, in flowering plants, such as *Oryza sativa* and *A. thaliana*, this linker has been implicated in binding the *alpha* subunit of the G protein heterotrimer (Pandey et al., 2009; Ma et al., 2015; Dong et al., 2019; Zhang et al., 2024).

To inspect whether certain polymorphisms are associated with major taxonomic groups, representative sequences of vascular plants and chordates were inspected without the gaps introduced by most unicellular organisms (Figure 3C). Notably, both mammals and flowering plants have clear consensus sequences respective to their own groups. For example, a member of an ancient Tracheophyte lineage (*Selaginella moellendorffii*) shows a variable region closely resembling that of flowering plants, while a distant chordate (*Petromyzon marinus)* displays a sequence that is closer to the one in mammals (Figure 3C). On the other hand, the choanoflagellate *Monosiga brevicollis*, which lacks some of the distinct insertions of unicellular organisms found in this region, shows substitutions that do not clearly align with the distinguishing characteristics of plants or animals (Figure 3C, black highlight). These analyses suggest that functional properties arising from this sequence segment, such as G_alpha_ binding, evolved within their respective taxonomic group.

### GPR89 can be classified as a Solute Carrier Protein

It has been suggested that GPR89 orthologs act either as a transporter or an ion channel (Maeda et al. 2009; Ma et al. 2015; Rojas et al. 2019). One of the most thorough functional studies was done in Trypanosoma brucei (Rojas et al. 2015), where authors suggest a role as an oligopeptide transporter, similar to SLC15 (Pfam PF00854), a member of the Major Facilitator Superfamily (MFS, CATH 1.20.1250.20) of solute carrier transporters (SLC). Members of the SLC superfamily are found throughout the eukaryotic cell in both the plasma membrane and organellar membranes. The SLC group comprises over 400 different transporters, organized in multiple families according to their sequence homology and transport characteristics (http://slc.bioparadigms.org/) (Tweedie et al., 2020; Ferrada and Superti-Furga, 2022).

Given the vast evolutionary distance between Trypanosoma and primates, which limits broader-scale comparisons, we investigated whether the human GPR89 ortholog is part of the large SLC transporter superfamily. SLC is a functional classification first defined by the HGNC (Hediger et al. 2004), whose members share very distant evidential homology at best. For this reason, we have decided to search for structural similarities, since protein structure is closely related to function, and is more sensitive to distant homologies than the primary sequence alone (Illergård et al., 2009; Chen et al., 2025). A preliminary search was performed using the predicted structure of human GPR89 (Uniprot ID: B7ZAQ6) against the full AlphaFold database via the Dali server (Varadi et al., 2022; Holm et al., 2023). Surprisingly, across the entire database, only the four members of the poorly studied LIMR family (Pfam PF04791) emerged as the closest structural matches (Table 1). The namesake member of this family, LIMR (i.e., LMBRL), has been associated with the endocytic transport of hydrophobic molecule-binding proteins, such as lipocalin-1 and β-lactoglobulin (Wojnar et al., 2003; Fluckinger et al., 2007). Another member, LMBRD1, participates in the lysosomal cobalamin transport, acting as an escort protein of ABCD4 transporter (Rutsch et al., 2009; Deme et al., 2014; Kawaguchi et al., 2016). Despite their reported indirect association to intracellular transport, no intrinsic transport activity has been demonstrated for these proteins. Further structural examination shows that they all share 9 transmembrane helices, with a linker between TM5 & 6 (Supplementary Figure 2).

**Table 1.** Structural comparison of human GPR89A against the AlphaFold Database using the Dali Server. Top hits (Z-scores > 10) are shown.

| Protein | Uniprot accession | %Id | RMSD | Z-score |
| --- | --- | --- | --- | --- |
| LMBD1 | Q9NUN5 | 15 | 5.6 | 19.6 |
| LMBD2 | Q68DH5 | 10 | 4.3 | 18.7 |
| LMBRL | Q6UX01 | 13 | 8.7 | 17.7 |
| LMBR1 | Q8WVP7 | 13 | 5.7 | 17.5 |

Next, we carried out an ambitious multiple structure alignment of over 480 human proteins, enabled by TM-Vec, a deep learning approach that efficiently approximates TM-scores without explicit structural superposition (Hamamsy et al., 2024). We included most human SLC and ABC transporters, plus representative members of other families, such as chlorine and calcium channels, and GPCRs. Structural outgroups were used to root the tree, i.e. beta-barrels (VDAC1 & TOM40) and globins (myo & hemoglobin). From the predicted TM-scores, we constructed a distance tree (1 - TM-score) using an unweighted pair group method with arithmetic mean (UPGMA) analysis (Figure 4). Despite employing a different analytical approach, our UPGMA tree largely replicated the “fold classification” groupings reported previously (Ferrada and Superti-Furga 2022) (Figure 4). Notably, our analysis revealed that the clade containing GPR89 is between the SLC39 (Zinc-Iron Permease family; Pfam PF02535, no CATH classification) and SLC42 (Ammonium transporters; Pfam PF00909, CATH superfamily 1.10.3430.10) clades (Figure 4B).

**Figure 4:**
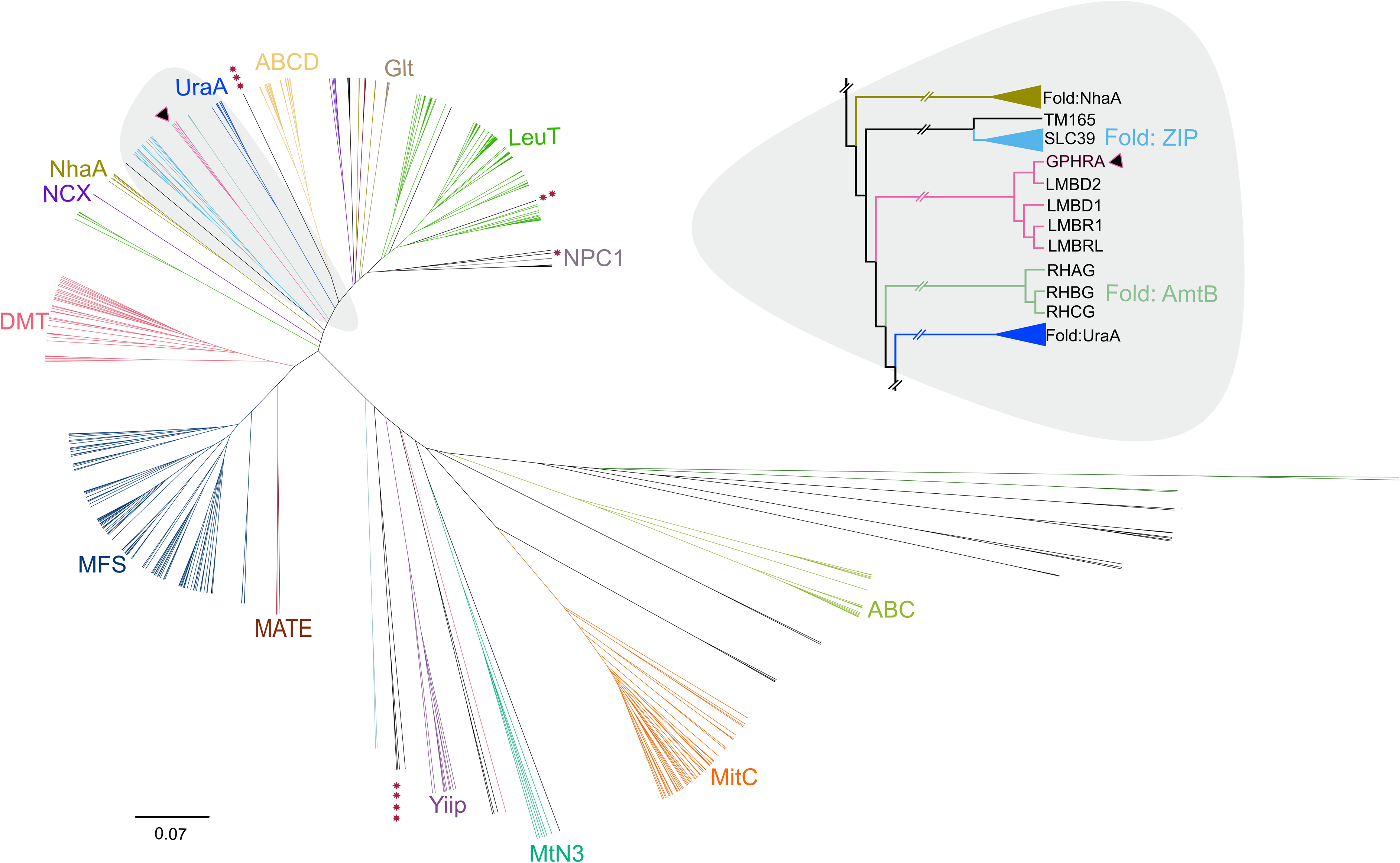
Unrooted UPGMA phylogenetic tree displays transporter protein sequences and human GPR89. **(A)** Expanded unrooted tree showing the SLC fold classification reported by Ferrada and Superti-Furga (2022) plus ABC, ABCD, CAC (*), CLCN (**), CFTR(***) and GPCR (*****) proteins. Branches are colored according to the SLC fold classification, thus black branches indicate other SLC proteins without fold classification. Black Triangle indicates GPHRA protein, and the gray highlight section is detailed in panel B. Dark green branches are the outgroup proteins. **(B)** Tree displaying cladogram representation focusing on clades relative to GRP89. Names of each protein and/or fold classification are shown on the tips.

In short, our evidence suggests that GPR89 is a novel member of the LIMR family and structurally related to the ZIP and AmtB families of SLC transporters.

### GPR89 shows a conserved transmembrane groove

To better understand the implications of ortholog similarities and differences, we inspected predicted structures available in the AlphaFold database (Varadi et al., 2022) for representative organisms of the various species groups previously mentioned. The predicted structure for human GPR89A (Uniprot ID B7ZAQ6), calculated by AlphaFold 2 (AF2) (Jumper et al., 2021; Varadi et al., 2022), shows the expected 9 transmembrane helices and a long cytosolic coiled-coil hairpin (HP) that shapes the TM5-TM6 linker (Figure 5). This is consistent with other predicted models in the database (e.g., Uniprot ID: *A. thaliana* (Q9XIP7), *D. discoideum* (Q54QM5), *T. brucei* (Q580H0); there is a systematic prediction of 9TM and a long helical hairpin forming the majority of the TM5-TM6 linker, likely constituting a structural signature for GPR89. In general, the large insertions displayed in various organisms occur predominantly in and around the cytosolic hairpin loop.

**Figure 5:**
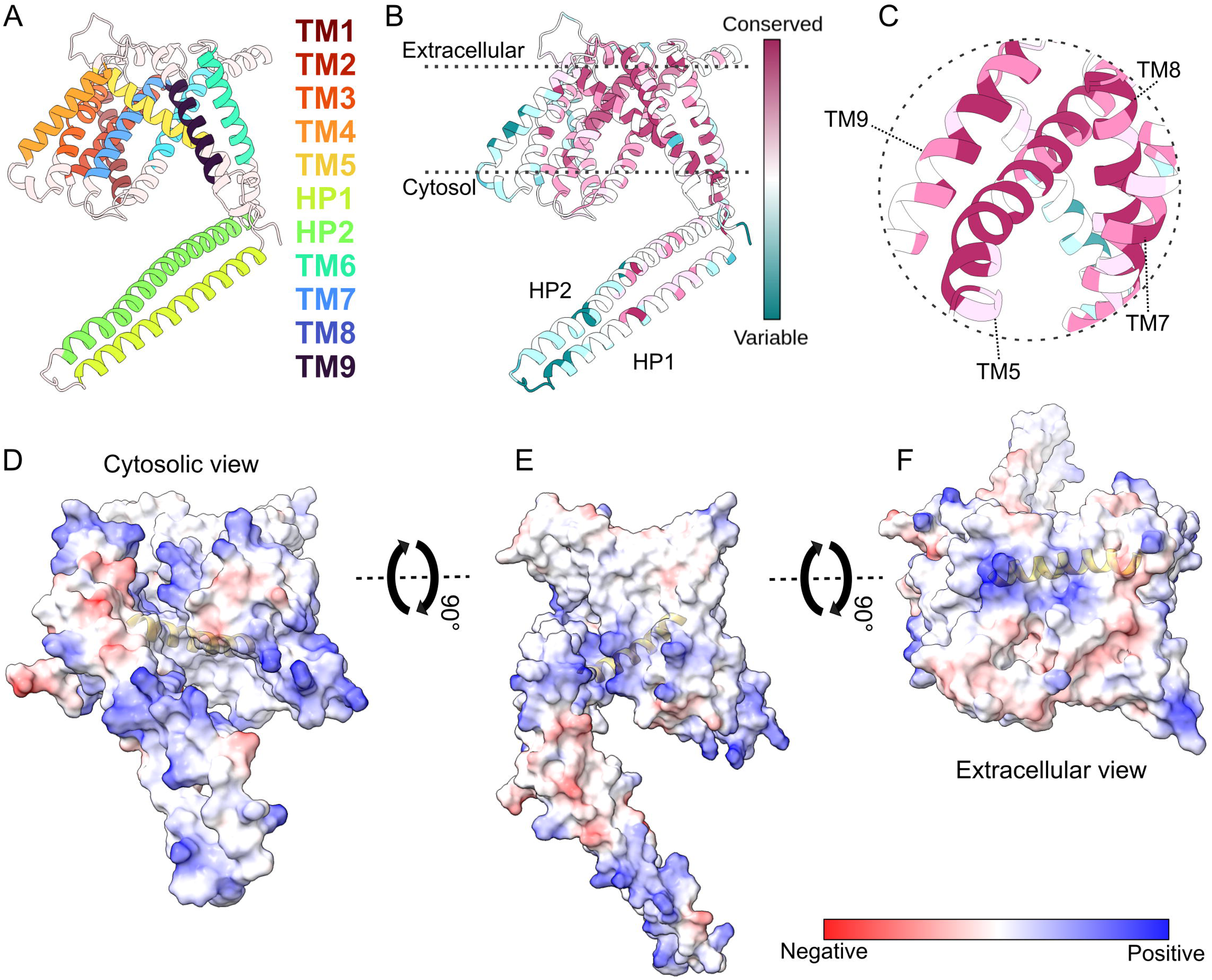
Predicted structure and conservation of human GPR89A. **(A)** AlphaFold2-predicted structure of GPR89A, showing 9 transmembrane helices and two cytosolic helices forming a hairpin. Colored TM residues correspond to those predicted by the DeepTMHMM server. **(B)** Conservation scores from ConSurf, using the MSA from figure 5, were mapped onto the structure and colored as a gradient from magenta (most) to cyan (least) conserved.**(C)** A closer look at the highly conserved TM5 and its vicinity. **(D-F)** Lateral hydrophilic cavity observed from the cytosolic **(D)**, membrane **(E)** and extracellular/luminal perspectives **(F)**. The surface is colored by calculated electrostatic potential; red: negative charge, blue: positive charge, white: neutral. TM5 is depicted in yellow.

To better visualize the relative position of the conserved residues, we displayed the conservation score at each position of the MSA in the context of the predicted structure for GPR89A (Figure 5B & C). By doing this structural mapping, we noted that conserved residues of different helices appear in close proximity (<4 Å atom to atom) to the well conserved TM5 helix, which appears tilted relative to the other transmembrane helices (Figure 5C). A large, partially hydrophilic vestibule, formed by TM5, TM6 & TM7, is accessible from the cytosolic face of the predicted structure (Figure 5D & E). This opening has a few charged and polar residues (R300, R357, Y390, H423, D427) that could form an aqueous pore. The cavity appears inaccessible from the extracellular side of the predicted structure (Figure 5F). It would be relevant to note that our model has not been optimized in explicit solvent and/or lipids.

Guided by the evidence that GPR89 is an SLC transporter, we modelled the protein together with putative ligands, i.e., small oligopeptides (Rojas et al., 2019). The new AlphaFold 3 server currently has a modest library of ligands to choose from, where di- or tripeptides are not available (Abramson et al., 2024). For this reason, We modelled human GPR89A with the tetrapeptide NHYG (Figure 6A-C), whose sequence was informed by the residues that produced the greatest physiological effect in *Trypanosoma brucei* (Rojas et al., 2019). The best model (ipTM = 0.65, pTM = 0.76) shows the peptide snugly fit into the lateral groove (Figure 6A-B), interacting mostly with polar residues, many of which have the maximum conservation score in the structure (Figure 6C). In the work of Rojas et al., the authors show that TbGPR89 transports the fluorescent oligopeptide analogue Ala-lys-AMCA. This ligand is available in an alternative deep learning modeller of protein-ligand structures: DynamicBind (Lu, et al., 2024). Here again, independently and without further user input, the oligopeptide is modelled inside the groove in not only the best model (lddt 0.449), but all 9 predicted models (Figure 6D-F & Supplementary data). In this case, however, the voluminous aromatic aminomethyl coumarin group is inserted deeply into the TM bundle, interacting with TM2 as well as TM5, 7 & 8 (Figure 6E). This prediction would be in good agreement with a putative transporter function.

**Figure 6.**
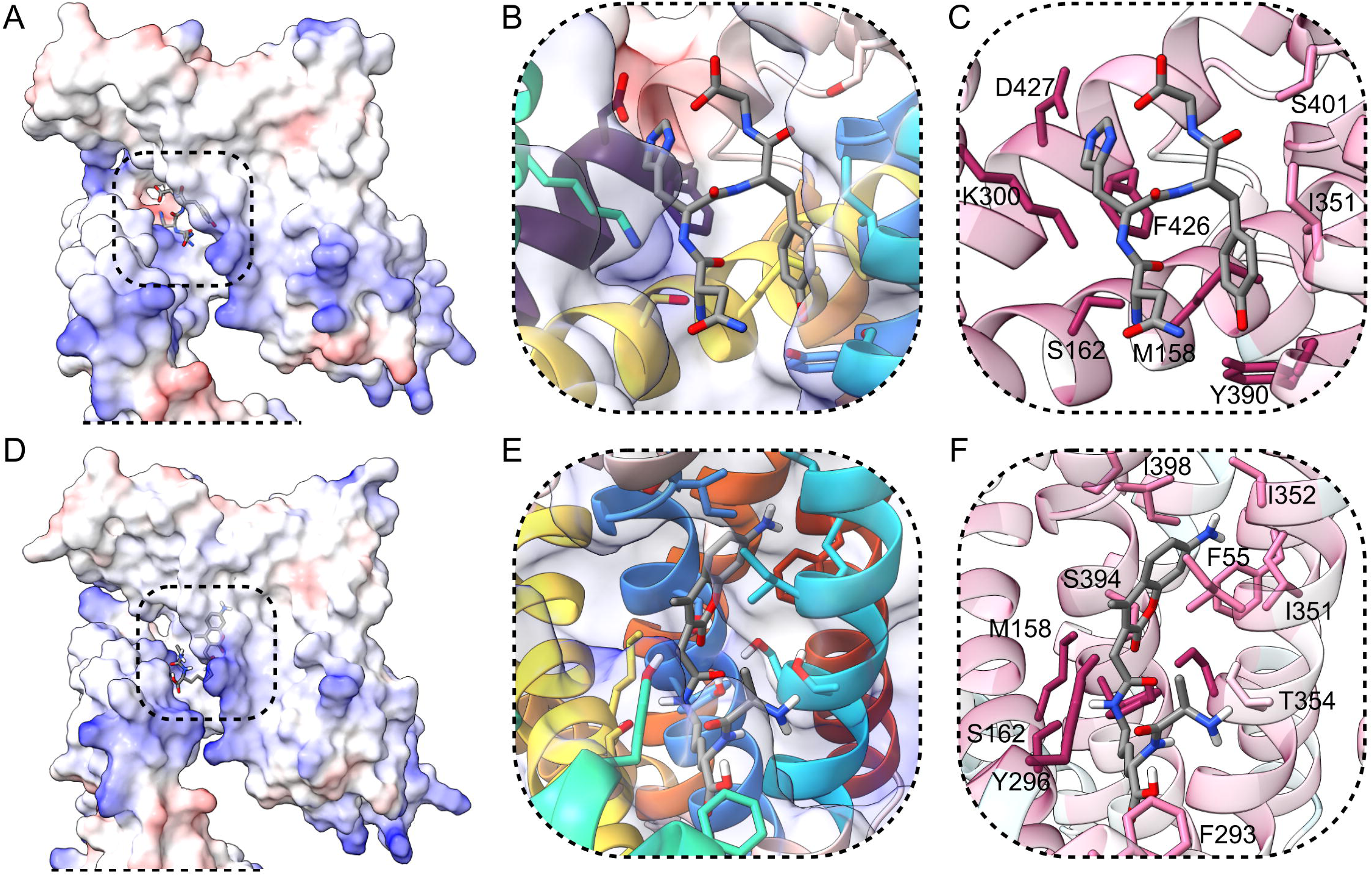
Peptide docking. **(A)** AlphaFold structural prediction of HsGPR89 with a small tetrapeptide. The groove is observed in detail in **(B & C). (D)** DynamicBind structural prediction of HsGPR89 with the fluorescent peptide analogue D-Ala-Lys-AMCA. The groove is observed in detail in **(E & F)**.

## DISCUSSION

GPR89 is widely conserved across eukaryotic lineages and is typically retained as a single-copy gene. Its robustness suggests a strong purifying selection and a function of general importance. At the same time, reports across taxa describe varied molecular roles and subcellular localizations. In vertebrates, GPR89 is found at the Golgi and endoplasmic reticulum membranes, where it likely supports the anion flux that is essential for maintaining luminal acidification. It has been suggested that GPR89 orthologs in plants mediate calcium permeability and contribute to the detection of hormonal and environmental cues. In *Trypanosoma brucei*, the protein localizes primarily to the plasma membrane where it supports the transport of small peptides. While this evidence may point to widespread functional divergence, an alternative explanation can be in the form of ion-coupled solute transport in which the preferred ion changes over evolutionary time. Our findings support the classification of GPR89 as a member of the solute carrier (SLC) superfamily, proposed originally by Rojas et al. (2019), with closer structural similarity to SLC39 and SLC42 families. The proposed classification is not based on novel biophysical data per se, but on the comparative structural and evolutionary analysis performed here. Notably, AlphaFold2-based structural models reveal a conserved transmembrane scaffold that is compatible with ion or solute translocation across membranes. In agreement with SLC classification (Bai et al., 2017; Pizzagalli et al., 2021), the structure displays a protein that is clearly separated in two sections and includes a central hydrophobic core surrounding a seemingly critical transmembrane segment (TM5) (Drew & Boudker, 2016; Colas et al., 2016). TM5 is consistently observed flanked by conserved transmembrane helices and polymorphic cytoplasmic and extracellular loops. These loops, that may be part of regulatory interactions or subcellular targeting signatures, display clade-specific sequence variation. This is particularly evident in the intracellular coiled-coil hairpin (HP) and the associated loop region (HPloop). Interestingly, the revealed lateral groove resembles that of the sodium-dependent bile salt transporter NTCP (Goutam et al., 2022; Liu et al., 2022), which is a close neighbor in the UPGMA structure tree. This architectural modularity would allow for the required physiological adaptation without altering the underlying transport capacity of the protein.

The phylogenetic distribution of GPR89 duplications provides additional evidence for its functional diversification. In vascular plants, multiple paralogs are often retained, sometimes with distinct expression patterns. In contrast, vertebrate GPR89 genes remain mostly single-copy, with evidence of pseudogenization in anthropoid primates. These lineage-specific events likely reflect regulatory specialization, as in regulating expression and hence the number of functional copies, rather than radical shifts in molecular function. Functional divergence through subcellular relocalization or necessary regulatory rewiring during organismal evolution could explain the differences in experimental observations, particularly in plants where conflicting localizations and signaling contexts have been reported.

While prior reports in plant systems have suggested GPCR activity and Gα protein binding (Pandey et al., 2009; Ma et al., 2015; Dong et al., 2019; Zhang et al., 2024), direct evidence for GPCR-like activation remains challenged, and no canonical downstream signaling pathway has been established. In contrast, the structural and phylogenetic context presented here supports a model in which GPR89 orthologs, as well as the sister group of LIMB proteins, might operate as transporters adapted to the specific demands of their subcellular environments.

Finally, the conservation of the core fold and its predicted compatibility with passive or facilitated transport suggest that evolutionary persistence shown by GPR89 is interlaced to its capacity to support fundamental aspects of organellar physiology, such as pH regulation in the Golgi, peptide uptake at the plasma membrane (likely ion-coupled), or ionic fluxes in the ER and endomembrane system. It would not be unreasonable to suggest that the variety of reported roles simply reflects changes in transported substrates, membrane context, and regulatory partners. This is not surprising in SLCs, where orthologous groups as well as related orphan genes sharing structural features, provide evidence of recent functional diversification and changes in transported substrates (Höglund *et al*., 2011; Ferrada and Superti-Furga 2022). Our perspective provides a framework to interpret observations that seem conflicting and underscores the need for further experimental work to define substrate specificity and transport kinetics.

In summary, our analysis places GPR89 within a well-supported structural and evolutionary context, consistent with a primary role in membrane transport. Phylogenetic stability, conserved transmembrane architecture, and classification within the human superfamily of SLC proteins suggests an ancient and adaptable transport function that has been carefully retained during eukaryotic evolution.

## MATERIALS AND METHODS

### Phylogenetic analyses

We used the ortholog prediction function of Ensembl database (Herrero et al., 2016) and NCBI BLAST (Altschul et al., 1990) to search for ortholog and paralog genes of *GPR89* on major groups of vertebrates, metazoan model species, major groups of Viridiplantae (including green algae), and unicellular eukaryotic organisms such as heterotrophic protists. After corroborating the number of transmembrane segments with DeepTMHMM and AlphaFold 2 DataBase (AF2-DB) (Hallgren et al., 2022; Varadi et al., 2022), the accuracy of the CDS translation was confirmed rendering an initial data set of 130 CDS including *GPR89A* and *GPR89B* copies for Archaeplastida, Heterotrophic protist, and Metazoa. (See details of the taxonomy sampling and the accessions number on Supplementary Table 1).

For the phylogenetic analyses on Supplementary Figure 1, we looked for orthologs and paralogs of *GPR89A* and *GPR89P* among major groups of Angiospermae and Mammals (Supplementary Table 2). In this case, we retrieved *GPR89P* performing a BLASTN search on the Ensembl database with default settings against the whole genome of model mammals organisms, using the functional CDS of each species as a query sequence. When *GRP89P* was present, we manually annotated the CDS using the coding exons of the functional gene of each species in Geneious 2023.2.1 (https://www.geneious.com). Therefore, we obtained two new data sets, the first one with 49 functional sequences from Angioespermae. The second contains 99 Mammalian sequences (including *GPR89A*, *B* and *P)*. *Amborella trichopoda* and Sauropsid sequences were used as outgroups for plants and animals respectively.

For all data sets, the Multiple Sequence Alignment (MSA) was performed using MAFFT (Katoh and Standley, 2013; Katoh et al., 2019) on Geneious Prime 2023.2.1, allowing the software to choose the strategy to determine best fit. Then, we performed a visual inspection of the alignments. The conservative sites were obtained on MEGA11 (Stecher et al., 2020), and Model Finder option was used by default on IQ-TREE 2 (Kalyaanamoorthy et al., 2017) to select the best fitting model for nucleotide substitutions (Figure 1: SYM+I+G4, Supplementary Figure 2A: GTR+F+I+G4, and 2B: SYM+I+G4). We further used IQ-TREE 2 (Minh et al., 2020) to conduct a maximum likelihood analysis in order to obtain the best phylogenetic tree. Bootstrapping for trees was performed to obtain the consensus from 1000 replicates. The analysis was run four times to ensure the accuracy. We report the best phylogenetic tree, obtained with the best Log-likelihood value (Figure 1: -105917.056, Supplementary Figure 2A: -21678.235, and 2B: -21013.558). The corresponding values for bootstrap support were obtained from the consensus tree. For details on Figure 1 and Figure 4 trees, you can find newick files on Supplementary material.

### Synteny assessment

From the Ensembl Compara database (Herrero et al., 2016) we obtained the position of the genes. From Genomicus 109.1 vertebrate database (Nguyen et al., 2022), and PLAZA5.0 Angiosperm database (Van Bel et al., 2022) we obtained the genomic context for predicted functional copies of GPR89. We examined upstream and downstream coding genes that are common to model organisms within these major groups. Specifically, for the case of mammals, we also look forward to retro-pseudogenes synteny but due to the fact that we were not able to find an evident pattern, synteny for pseudogenes is not reported.

### Amino acidic frequency analysis

The 128 non-redundant sequences from the CDS were translated *in silico*, and then the amino acid sequences were aligned using MAFFT (L-INS-i strategy). In order to facilitate the visualization in Figure 3C, we minimized the number and length of the gaps by removing protists and other sequences, without realignment.

### Multiple Structural Alignment

We selected a collection of human transporters, defined as such in the IUPHAR/BPS Guide to Pharmacology (Alexander et al., 2023). Sequences included the Solute Carriers (SLC) reported in Ferrada and Superti-Furga, 2022; ABC transporters, human GPR89A, LMB protein homologs and representatives of other protein families. Beta-barrel proteins (VDAC1 and TOM40) & globulins (myoglobin and alpha-hemoglobin) were chosen as structural outgroups. The amino acid sequence of these proteins was obtained directly from Uniprot, via an automatic Python script and manual accession. Transporter subunit OSTB was excluded from the analysis for its short length (128 a.a.). The final list comprises 482 sequences in total.

The structural alignment was carried out using TM-Vec (Hamamsy et al., 2024). This Deep-Learning tool predicts the TM-score (structural similarity) of proteins via their primary structure. We used the large CATH dataset-trained model (Hamamsy et al., 2024) and the full amino acid sequences previously described. The TM-score prediction was turned to a symmetric distance matrix, where structural distance was defined as 1 - TM-score (0 = identical structures, 1 = complete dissimilar structures). Then we ran the UPGMA analysis on MAFFT online (https://mafft.cbrc.jp/alignment/server/phylogeny.html). We used the unrooted ultrametric tree. The tree topology was then compared and annotated according to the results reported by Ferrada and Superti-Furga, 2022. This tree file can be found in newick format on our supplementary material S2.

### Structural predictions

All 3D models for the different GPR89 orthologs and paralogs we found present in Eukarya were obtained from the freely available AlphaFold-DB (Varadi et al., 2022). We present human GPR89A as a representative structural model. All inspected models shared the general 9TM topology and the structural signatures discussed in this paper.

Protein-peptide dockings were predicted using the AlphaFold3 server & DynamicBind v2 (Lu et al., 2024; Abramson et al., 2024).

### Figure rendering and accessibility

The trees were visualized using Figtree 1.4.4 (http://tree.bio.ed.ac.uk/software/figtree/) and vectorial silhouettes for phylogenetic figures were taken from https:/phylopic.org. These figures were colored and prepared on Inkscape 1.1.2.

MSA data was plotted with a custom Python script (Cabezas-Bratesco et al., 2022) and the sequence logo was made using LogoMaker (Tareen & Kinney, 2020). For accessibility, Google’s Turbo color palette was chosen for labeling (http://research.google/blog/turbo-an-improved-rainbow-colormap-for-visualization/), and all figures were tested using a Color Blindness Simulator in Python (https://medium.com/@er_95882/colour-blindness-simulator-in-python-detecting-visualisation-problems-in-presentations-and-reports-71595526dc0e).

The degree of conservation of the different residues obtained from the MSA was mapped onto the structure of GPR89 using the web server ConSurf (Glaser et al., 2003; Ashkenazy et al., 2016). The mapped structure was then exported to the molecular visualization software UCSF ChimeraX (Goddard et al., 2018; Pettersen et al., 2021), and the conservation score was manually annotated into the PDB file. The transmembrane helices were annotated according to the prediction of the DeepTMHMM server (Hallgren et al., 2022). Coulombic surface potentials were calculated by ChimeraX.

## Acknowledgements.

The authors acknowledge Dr. Juan C. Opazo for consultation on annotating pseudogene sequences. This work was supported by Beca ANID #21220073 to LR, Fondo Nacional de Desarrollo Científico y Tecnológico from the Chilean government (FONDECYT 1191868 and 1241753) to SB, and ANID-Millennium Science Initiative Program #NCN19_168 (SB and CQ). The Millennium Nucleus of Ion Channel-Associated Diseases (MiNICAD) is a Millennium Nucleus of the Iniciativa Milenio, National Agency of Research and Development (ANID, Chile). This research was supported by the *Patagon* supercomputer of *Universidad Austral de Chile* (FONDEQUIP EQM180042).

## Author contributions

Camila A. Quercia-Raty, Formal analysis, Data curation, Investigation, Visualization, Writing – original draft.

Luka Robeson, Formal analysis, Methodology, Visualization, Writing – review and editing.

Sebastian E Brauchi, Conceptualization, Data curation, Formal analysis, Methodology, Project administration, Writing – original draft, Writing – review and editing.

## Declaration of generative AI and AI-assisted technologies in the writing process

During the preparation of this work, the authors used ChatGPT by OpenAI to correct the language and improve the readability of the text. After using this tool, the authors reviewed and edited the content as needed and take full responsibility for the content of the published article.

## REFERENCE LIST

Abramson, J., Adler, J., Dunger, J. et al. (2024). Accurate structure prediction of biomolecular interactions with AlphaFold 3. Nature 630, 493–500. doi: 10.1038/s41586-024-07487-w

Albalat, R., and Cañestro, C. (2016). Evolution by gene loss. Nat. Rev. Genet. 17, 379–391. doi: 10.1038/nrg.2016.39

Alexander, S. P., Fabbro, D., Kelly, E., Mathie, A. A., Peters, J. A., Veale, E. L., et al. (2023). The concise guide to PHARMACOLOGY 2023/24: transporters. British Journal of Pharmacology, 180, S374–S469. doi: 10.1111/bph.16182

Altschul, S. F., Gish, W., Miller, W., Myers, E. W., and Lipman, D. J. (1990). Basic local alignment search tool. J. Mol. Biol. 215, 403–410. doi: 10.1016/S0022-2836(05)80360-2

Ashkenazy, H., Abadi, S., Martz, E., Chay, O., Mayrose, I., Pupko, T., et al. (2016). ConSurf 2016: an improved methodology to estimate and visualize evolutionary conservation in macromolecules. Nucleic Acids Res. 44, W344–W350. doi: 10.1093/nar/gkw408

Bai, X., Moraes, T. F., and Reithmeier, R. A. F. (2017). Structural biology of solute carrier (SLC) membrane transport proteins. Mol. Membr. Biol. 34, 1–32. doi: 10.1080/09687688.2018.1448123

Bergsten, J. (2005). A review of long-branch attraction. Cladistics 21, 163–193. doi: 10.1111/j.1096-0031.2005.00059.x

Burki, F., Roger, A. J., Brown, M. W., and Simpson, A. G. B. (2020). The New Tree of Eukaryotes. Trends Ecol. Evol. 35, 43–55. doi: 10.1016/j.tree.2019.08.008

Cabezas-Bratesco, D., Mcgee, F. A., Colenso, C. K., Zavala, K., Granata, D., Carnevale, V., et al. (2022). Sequence and structural conservation reveal fingerprint residues in TRP channels. eLife 11, e73645. doi: 10.7554/eLife.73645

Calebiro, D., Koszegi, Z., Lanoiselée, Y., Miljus, T., and O’Brien, S. (2021). G protein-coupled receptor-G protein interactions: a single-molecule perspective. Physiol. Rev. 101, 857– 906. doi: 10.1152/physrev.00021.2020

Cañestro, C., Albalat, R., Irimia, M., and Garcia-Fernàndez, J. (2013). Impact of gene gains, losses and duplication modes on the origin and diversification of vertebrates. Semin. Cell Dev. Biol. 24, 83–94. doi: 10.1016/j.semcdb.2012.12.008

Charroux, B., and Royet, J. (2013). Mutations in the Drosophila ortholog of the vertebrate Golgi pH regulator (GPHR) protein disturb endoplasmic reticulum and Golgi organization and affect systemic growth. Biol. Open 3, 72–80. doi: 10.1242/bio.20137187

Chen, Z., Zhang, X., Yu, W., & Luo, Q. (2025). A comprehensive benchmarking study of protein structure alignment tools based on downstream task performance. bioRxiv, 2025-03. doi: 10.1101/2025.03.11.642719

Cheng, F., Wu, J., Cai, X., Liang, J., Freeling, M., and Wang, X. (2018). Gene retention, fractionation and subgenome differences in polyploid plants. Nature Plants 4, 258–268. doi: 10.1038/s41477-018-0136-7

Ciomborowska-Basheer, J., Staszak, K., Kubiak, M. R., and Makałowska, I. (2021). Not So Dead Genes—Retrocopies as Regulators of Their Disease-Related Progenitors and Hosts. Cells 10, 912. doi: 10.3390/cells10040912

Colas, C., Ung, P. M. U., & Schlessinger, A. (2016). SLC transporters: structure, function, and drug discovery. Medchemcomm, 7(6), 1069–1081. doi: 10.1039/C6MD00005C

Cooper, G. M. (2000). “The Golgi Apparatus,” in The Cell: A Molecular Approach. 2nd edition, (Sinauer Associates). Available at: https://www.ncbi.nlm.nih.gov/books/NBK9838/ (Accessed October 5, 2023).

Deckstein, J., van Appeldorn, J., Tsangarides, M., Yiannakou, K., Müller, R., Stumpf, M., et al. (2015). The Dictyostelium discoideum GPHR Ortholog Is an Endoplasmic Reticulum and Golgi Protein with Roles during Development. Eukaryot. Cell 14, 41–54. doi: 10.1128/EC.00208-14

Deme, J. C., Hancock, M. A., Xia, X., Shintre, C. A., Plesa, M., Kim, J. C., et al. (2014). Purification and interaction analyses of two human lysosomal vitamin B_12_ transporters: LMBD1 and ABCD4. Molecular Membrane Biology, 31(7–8), 250–261. doi: 10.3109/09687688.2014.990998

Dennis, M. Y., Harshman, L., Nelson, B. J., Penn, O., Cantsilieris, S., Huddleston, J., et al. (2017). The evolution and population diversity of human-specific segmental duplications. *Nat*. Ecol. Evol. 1, 0069. doi: 10.1038/s41559-016-0069

Dong, H., Yan, S., Liu, J., Liu, P., and Sun, J. (2019). TaCOLD1 defines a new regulator of plant height in bread wheat. Plant Biotechnol. J. 17, 687–699. doi: 10.1111/pbi.13008

Drew, D., and Boudker, O. (2016). Shared Molecular Mechanisms of Membrane Transporters. Annu. Rev. Biochem. 85, 543–572. doi: 10.1146/annurev-biochem-060815-014520

Felsenstein, J. (1978). Cases in which Parsimony or Compatibility Methods Will be Positively Misleading. Syst. Zool. 27, 401–410. doi: 10.2307/2412923

Ferrada, E., and Superti-Furga, G. (2022). A structure and evolutionary-based classification of solute carriers. iScience 25, 105096. doi: 10.1016/j.isci.2022.105096

Fluckinger, M., Merschak, P., Hermann, M., Haertlé, T., & Redl, B. (2008). Lipocalin-interacting-membrane-receptor (LIMR) mediates cellular internalization of β-lactoglobulin. Biochimica et Biophysica Acta (BBA)-Biomembranes, 1778(1), 342–347. doi: 10.1016/j.bbamem.2007.10.010

García-Nafría, J., and Tate, C. G. (2019). Cryo-EM structures of GPCRs coupled to Gs, Gi and Go. Mol. Cell. Endocrinol. 488, 1–13. doi: 10.1016/j.mce.2019.02.006

Glaser, F., Pupko, T., Paz, I., Bell, R. E., Bechor-Shental, D., Martz, E., et al. (2003). ConSurf: Identification of Functional Regions in Proteins by Surface-Mapping of Phylogenetic Information. Bioinformatics 19, 163–164. doi: 10.1093/bioinformatics/19.1.163

Goddard, T. D., Huang, C. C., Meng, E. C., Pettersen, E. F., Couch, G. S., Morris, J. H., et al. (2018). UCSF ChimeraX: Meeting modern challenges in visualization and analysis. Protein Sci. Publ. Protein Soc. 27, 14–25. doi: 10.1002/pro.3235

Goutam, K., Ielasi, F.S., Pardon, E. et al. (2022). Structural basis of sodium-dependent bile salt uptake into the liver. Nature 606, 1015–1020. doi: 10.1038/s41586-022-04723-z

Hallgren, J., Tsirigos, K. D., Pedersen, M. D., Armenteros, J. J. A., Marcatili, P., Nielsen, H., et al. (2022). DeepTMHMM predicts alpha and beta transmembrane proteins using deep neural networks. bioRxiv 2022.04.08.487609. doi: 10.1101/2022.04.08.487609

Hamamsy, T., Morton, J.T., Blackwell, R. et al. (2024). Protein remote homology detection and structural alignment using deep learning. Nat Biotechnol 42, 975–985. doi: 10.1038/s41587-023-01917-2

Hediger, M.A., Romero, M.F., Peng, JB. et al. (2004). The ABCs of solute carriers: physiological, pathological and therapeutic implications of human membrane transport proteins. Pflugers Arch - Eur J Physiol 447, 465–468. doi: 10.1007/s00424-003-1192-y

Herrero, J., Muffato, M., Beal, K., Fitzgerald, S., Gordon, L., Pignatelli, M., et al. (2016). Ensembl comparative genomics resources. Database 2016, baw053. doi: 10.1093/database/baw053

Heslop-Harrison, J. S., Schwarzacher, T., and Liu, Q. (2023). Polyploidy: Its consequences and enabling role in plant diversification and evolution. Annals of Botany 131, 1–10. doi: 10.1093/aob/mcac132

Hilger, D., Masureel, M., and Kobilka, B. K. (2018). Structure and dynamics of GPCR signaling complexes. Nat. Struct. Mol. Biol. 25, 4–12. doi: 10.1038/s41594-017-0011-7

Holm, L., Laiho, A., Törönen, P., & Salgado, M. (2023). DALI shines a light on remote homologs: One hundred discoveries. Protein Science, 32(1), e4519. doi: 10.1002/pro.4519

Hopf, T. A., Colwell, L. J., Sheridan, R., Rost, B., Sander, C., and Marks, D. S. (2012). Three-Dimensional Structures of Membrane Proteins from Genomic Sequencing. Cell 149, 1607–1621. doi: 10.1016/j.cell.2012.04.012

Hopf, T. A., Ingraham, J. B., Poelwijk, F. J., Schärfe, C. P. I., Springer, M., Sander, C., et al. (2017). Mutation effects predicted from sequence co-variation. Nat. Biotechnol. 35, 128–135. doi: 10.1038/nbt.3769

Höglund, P. J., Nordström, K. J., Schiöth, H. B., & Fredriksson, R. (2011). The solute carrier families have a remarkably long evolutionary history with the majority of the human families present before divergence of Bilaterian species. Molecular biology and evolution, 28(4), 1531–1541. doi:

Illergård, K., Ardell, D. H., & Elofsson, A. (2009). Structure is three to ten times more conserved than sequence—a study of structural response in protein cores. *Proteins: Structure*, Function, and Bioinformatics, 77(3), 499–508. doi: 10.1002/prot.22458

Jaffé, F. W., Freschet, G.-E. C., Valdes, B. M., Runions, J., Terry, M. J., and Williams, L. E. (2012). G Protein–Coupled Receptor-Type G Proteins Are Required for Light-Dependent Seedling Growth and Fertility in Arabidopsis. Plant Cell 24, 3649–3668. doi: 10.1105/tpc.112.098681

Jentsch, T. J., and Pusch, M. (2018). CLC Chloride Channels and Transporters: Structure, Function, Physiology, and Disease. Physiol. Rev. 98, 1493–1590. doi: 10.1152/physrev.00047.2017

Jumper, J., Evans, R., Pritzel, A., Green, T., Figurnov, M., Ronneberger, O., et al. (2021). Highly accurate protein structure prediction with AlphaFold. Nature 596, 583–589. doi: 10.1038/s41586-021-03819-2

Kalyaanamoorthy, S., Minh, B. Q., Wong, T. K. F., von Haeseler, A., and Jermiin, L. S. (2017). ModelFinder: fast model selection for accurate phylogenetic estimates. Nat. Methods 14, 587–589. doi: 10.1038/nmeth.4285

Katoh, K., Rozewicki, J., and Yamada, K. D. (2019). MAFFT online service: multiple sequence alignment, interactive sequence choice and visualization. Brief. Bioinform. 20, 1160– 1166. doi: 10.1093/bib/bbx108

Katoh, K., and Standley, D. M. (2013). MAFFT Multiple Sequence Alignment Software Version 7: Improvements in Performance and Usability. Mol. Biol. Evol. 30, 772–780. doi: 10.1093/molbev/mst010

Kawaguchi, K., Okamoto, T., Morita, M. et al. (2016). Translocation of the ABC transporter ABCD4 from the endoplasmic reticulum to lysosomes requires the escort protein LMBD1. Sci Rep 6, 30183. doi: 10.1038/srep30183

Khosrowabadi, E., & Kellokumpu, S. (2020). Golgi pH and ion homeostasis in health and disease. Organelles in Disease, 1–23. doi: 10.1007/112_2020_49

Kornak, U., Reynders, E., Dimopoulou, A., van Reeuwijk, J., Fischer, B., Rajab, A., et al. (2008). Impaired glycosylation and cutis laxa caused by mutations in the vesicular H+-ATPase subunit ATP6V0A2. Nat. Genet. 40, 32–34. doi: 10.1038/ng.2007.45

Kumar, S., Suleski, M., Craig, J. M., Kasprowicz, A. E., Sanderford, M., Li, M., et al. (2022). TimeTree 5: An Expanded Resource for Species Divergence Times. Molecular Biology and Evolution 39, msac174. doi: 10.1093/molbev/msac174

Kunzelmann, K. (2015). TMEM16, LRRC8A, bestrophin: chloride channels controlled by Ca2+ and cell volume. Trends Biochem. Sci. 40, 535–543. doi: 10.1016/j.tibs.2015.07.005

Kuzmin, E., Taylor, J. S., and Boone, C. (2022). Retention of duplicated genes in evolution. Trends in Genetics 38, 59–72. doi: 10.1016/j.tig.2021.06.016

Levy, A. A., and Feldman, M. (2022). Evolution and origin of bread wheat. Plant Cell 34, 2549– 2567. doi: 10.1093/plcell/koac130

Li, J., and Wang, Y. (2022). Golgi Metal Ion Homeostasis in Human Health and Diseases. Cells 11, 289. doi: 10.3390/cells11020289

Liu, C., Wu, Y., Liu, Y., Yang, L., Dong, R., Jiang, L., et al. (2021). Genome-wide analysis of tandem duplicated genes and their contribution to stress resistance in pigeonpea (Cajanus cajan). Genomics 113, 728–735. doi: 10.1016/j.ygeno.2020.10.003

Liu, F., Zhang, Z., Csanády, L., Gadsby, D. C., and Chen, J. (2017). Molecular Structure of the Human CFTR Ion Channel. Cell 169, 85–95.e8. doi: 10.1016/j.cell.2017.02.024

Liu, H., Irobalieva, R.N., Bang-Sørensen, R. et al. (2022). Structure of human NTCP reveals the basis of recognition and sodium-driven transport of bile salts into the liver. Cell Res 32, 773–776. doi: 10.1038/s41422-022-00680-4

Lu, W., Zhang, J., Huang, W. et al. (2024). DynamicBind: predicting ligand-specific protein-ligand complex structure with a deep equivariant generative model. Nat Commun 15, 1071. doi: 10.1038/s41467-024-45461-2

Ma, Y., Dai, X., Xu, Y., Luo, W., Zheng, X., Zeng, D., et al. (2015). COLD1 Confers Chilling Tolerance in Rice. Cell 160, 1209–1221. doi: 10.1016/j.cell.2015.01.046

Maeda, Y., Ide, T., Koike, M., Uchiyama, Y., and Kinoshita, T. (2008). GPHR is a novel anion channel critical for acidification and functions of the Golgi apparatus. Nat. Cell Biol. 10, 1135–1145. doi: 10.1038/ncb1773

Marks, D. S., Colwell, L. J., Sheridan, R., Hopf, T. A., Pagnani, A., Zecchina, R., et al. (2011). Protein 3D Structure Computed from Evolutionary Sequence Variation. PLOS ONE 6, e28766. doi: 10.1371/journal.pone.0028766

Mighell, A. J., Smith, N. R., Robinson, P. A., and Markham, A. F. (2000). Vertebrate pseudogenes. FEBS Lett. 468, 109–114. doi: 10.1016/s0014-5793(00)01199-6

Minh, B. Q., Schmidt, H. A., Chernomor, O., Schrempf, D., Woodhams, M. D., von Haeseler, A., et al. (2020). IQ-TREE 2: New Models and Efficient Methods for Phylogenetic Inference in the Genomic Era. Mol. Biol. Evol. 37, 1530–1534. doi: 10.1093/molbev/msaa015

Mojib, N., and Kubanek, J. (2020). Comparative transcriptomics supports the presence of G protein-coupled receptor-based signaling in unicellular marine eukaryotes. Limnol. Oceanogr. 65, 762–774. doi: 10.1002/lno.11345

Morava, É., Guillard, M., Lefeber, D. J., and Wevers, R. A. (2009). Autosomal recessive cutis laxa syndrome revisited. Eur. J. Hum. Genet. 17, 1099–1110. doi: 10.1038/ejhg.2009.22

Nguyen, N. T. T., Vincens, P., Dufayard, J. F., Roest Crollius, H., and Louis, A. (2022). Genomicus in 2022: comparative tools for thousands of genomes and reconstructed ancestors. Nucleic Acids Res. 50, D1025–D1031. doi: 10.1093/nar/gkab1091

Nordeen, M. H., Jones, S. M., Howell, K. E., and Caldwell, J. H. (2000). GOLAC: an endogenous anion channel of the Golgi complex. Biophys. J. 78, 2918–2928. doi: 10.1016/S0006-3495(00)76832-9

Omasits, U., Ahrens, C. H., Müller, S., & Wollscheid, B. (2014). Protter: interactive protein feature visualization and integration with experimental proteomic data. Bioinformatics, 30(6), 884–886.

Osei-Owusu, J., Yang, J., Vitery, M. D. C., and Qiu, Z. (2018). Molecular Biology and Physiology of Volume-Regulated Anion Channel (VRAC). Curr. Top. Membr. 81, 177–203. doi: 10.1016/bs.ctm.2018.07.005

Panchy, N., Lehti-Shiu, M., and Shiu, S.-H. (2016). Evolution of Gene Duplication in Plants. Plant Physiology 171, 2294–2316. doi: 10.1104/pp.16.00523

Pandey, S., Nelson, D. C., and Assmann, S. M. (2009). Two Novel GPCR-Type G Proteins Are Abscisic Acid Receptors in Arabidopsis. Cell 136, 136–148. doi: 10.1016/j.cell.2008.12.026

Paroutis, P., Touret, N., and Grinstein, S. (2004). The pH of the Secretory Pathway: Measurement, Determinants, and Regulation. Physiology 19, 207–215. doi: 10.1152/physiol.00005.2004

Pettersen, E. F., Goddard, T. D., Huang, C. C., Meng, E. C., Couch, G. S., Croll, T. I., et al. (2021). UCSF ChimeraX: Structure visualization for researchers, educators, and developers. Protein Sci. Publ. Protein Soc. 30, 70–82. doi: 10.1002/pro.3943

Pizzagalli, M. D., Bensimon, A., and Superti-Furga, G. (2021). A guide to plasma membrane solute carrier proteins. FEBS J. 288, 2784–2835. doi: 10.1111/febs.15531

Ren, R., Wang, H., Guo, C., Zhang, N., Zeng, L., Chen, Y., Ma, H., and Qi, J. (2018). Widespread Whole Genome Duplications Contribute to Genome Complexity and Species Diversity in Angiosperms. Molecular Plant 11, 414–428. doi: 10.1016/j.molp.2018.01.002

Rivinoja, A., Kokkonen, N., Kellokumpu, I., and Kellokumpu, S. (2006). Elevated Golgi pH in breast and colorectal cancer cells correlates with the expression of oncofetal carbohydrate T-antigen. J. Cell. Physiol. 208, 167–174. doi: 10.1002/jcp.20653

Rivinoja, A., Pujol, F. M., Hassinen, A., and Kellokumpu, S. (2012). Golgi pH, its regulation and roles in human disease. Ann. Med. 44, 542–554. doi: 10.3109/07853890.2011.579150

Rojas, F., Silvester, E., Young, J., Milne, R., Tettey, M., Houston, D. R., et al. (2019). Oligopeptide Signaling through TbGPR89 Drives Trypanosome Quorum Sensing. Cell 176, 306–317.e16. doi: 10.1016/j.cell.2018.10.041

Rutsch, F., Gailus, S., Miousse, I. et al. (2009). Identification of a putative lysosomal cobalamin exporter altered in the cblF defect of vitamin B_12_ metabolism. Nat Genet 41, 234–239. doi: 10.1038/ng.294

Sabirov, R. Z., Merzlyak, P. G., Islam, M. R., Okada, T., and Okada, Y. (2016). The properties, functions, and pathophysiology of maxi-anion channels. Pflugers Arch. 468, 405–420. doi: 10.1007/s00424-015-1774-5

Sabirov, R. Z., Merzlyak, P. G., Okada, T., Islam, M. R., Uramoto, H., Mori, T., et al. (2017). The organic anion transporter SLCO2A1 constitutes the core component of the Maxi-Cl channel. EMBO J. 36, 3309–3324. doi: 10.15252/embj.201796685

Sato, Y., and Nishida, M. (2010). Teleost fish with specific genome duplication as unique models of vertebrate evolution. Environmental Biology of Fishes 88, 169–188. doi:10.1007/s10641-010-9628-7

Sauguet, L., Shahsavar, A., and Delarue, M. (2015). Crystallographic studies of pharmacological sites in pentameric ligand-gated ion channels. Biochim. Biophys. Acta BBA - Gen. Subj. 1850, 511–523. doi: 10.1016/j.bbagen.2014.05.007

Shiu, S. H., Byrnes, J. K., Pan, R., Zhang, P., and Li, W. H. (2006). Role of positive selection in the retention of duplicate genes in mammalian genomes. Proc. Natl. Acad. Sci. 103, 2232–2236. doi: 10.1073/pnas.0510388103

Soltis, P. S., and Soltis, D. E. (2016). Ancient WGD events as drivers of key innovations in angiosperms. Curr. Opin. Plant Biol. 30, 159–165. doi: 10.1016/j.pbi.2016.03.015

Sou, Y., Kakuta, S., Kamikubo, Y., Niisato, K., Sakurai, T., Parajuli, L. K., et al. (2019). Cerebellar Neurodegeneration and Neuronal Circuit Remodeling in Golgi pH Regulator-Deficient Mice. eNeuro 6, ENEURO.0427-18.2019. doi: 10.1523/ENEURO.0427-18.2019

Sou, Y., Yamaguchi, J., Kameda, H., Masuda, K., Maeda, Y., Uchiyama, Y., et al. (2022). GPHR-mediated acidification of the Golgi lumen is essential for cholesterol biosynthesis in the brain. FEBS Lett. 596, 2873–2888. doi: 10.1002/1873-3468.14491

Spano, D., and Colanzi, A. (2022). Golgi Complex: A Signaling Hub in Cancer. Cells 11, 1990. doi: 10.3390/cells11131990

Stecher, G., Tamura, K., and Kumar, S. (2020). Molecular Evolutionary Genetics Analysis (MEGA) for macOS. Molecular Biology and Evolution 37, 1237–1239. doi:10.1093/molbev/msz312

Syrovatkina, V., Alegre, K. O., Dey, R., and Huang, X.-Y. (2016). Regulation, Signaling, and Physiological Functions of G-Proteins. J. Mol. Biol. 428, 3850–3868. doi: 10.1016/j.jmb.2016.08.002

Tareen, A., & Kinney, J. B. (2020). Logomaker: Beautiful sequence logos in Python. Bioinformatics 36, 2272–2274. doi:10.1093/bioinformatics/btz921

Tarutani, M., Nakajima, K., Uchida, Y., Takaishi, M., Goto-Inoue, N., Ikawa, M., et al. (2012). GPHR-Dependent Functions of the Golgi Apparatus Are Essential for the Formation of Lamellar Granules and the Skin Barrier. J. Invest. Dermatol. 132, 2019–2025. doi: 10.1038/jid.2012.100

The Angiosperm Phylogeny Group. (2016). An update of the Angiosperm Phylogeny Group classification for the orders and families of flowering plants: APG IV. Botanical Journal of the Linnean Society 181, 1–20. doi:10.1111/boj.12385

Thompson, R. J., Nordeen, M. H., Howell, K. E., and Caldwell, J. H. (2002). A Large-Conductance Anion Channel of the Golgi Complex. Biophys. J. 83, 278–289. doi: 10.1016/S0006-3495(02)75168-0

Tweedie, S., Braschi, B., Gray, K., Jones, T. E. M., Seal, R. L., Yates, B., et al. (2020). Genenames.org: the HGNC and VGNC resources in 2021. Nucleic Acids Res. 49, D939–D946. doi: 10.1093/nar/gkaa980

Van Bel, M., Silvestri, F., Weitz, E. M., Kreft, L., Botzki, A., Coppens, F., et al. (2022). PLAZA 5.0: Extending the scope and power of comparative and functional genomics in plants. Nucleic Acids Research 50, D1468–D1474. doi:10.1093/nar/gkab1024

Van De Peer, Y., Mizrachi, E., and Marchal, K. (2017). The evolutionary significance of polyploidy. Nature Reviews Genetics 18, 411–424. doi:10.1038/nrg.2017.26

Varadi, M., Anyango, S., Deshpande, M., Nair, S., Natassia, C., Yordanova, G., et al. (2022). AlphaFold Protein Structure Database: massively expanding the structural coverage of protein-sequence space with high-accuracy models. Nucleic Acids Res. 50, D439–D444. doi: 10.1093/nar/gkab1061

Very, N., Lefebvre, T., and Yazidi-Belkoura, I. E. (2017). Drug resistance related to aberrant glycosylation in colorectal cancer. Oncotarget 9, 1380–1402. doi: 10.18632/oncotarget.22377

Wei, N., Cronn, R., Liston, A., and Ashman, T. (2019). Functional trait divergence and trait plasticity confer polyploid advantage in heterogeneous environments. New Phytologist 221, 2286–2297. doi: 10.1111/nph.15508

Wojnar, P., Lechner, M., & Redl, B. (2003). Antisense down-regulation of lipocalin-interacting membrane receptor expression inhibits cellular internalization of lipocalin-1 in human NT2 cells. Journal of Biological Chemistry, 278(18), 16209–16215. doi: 10.1074/jbc.M210922200

Zhang, J. (2003). Evolution by gene duplication: an update. Trends Ecol. Evol. 18, 292–298. doi: 10.1016/S0169-5347(03)00033-8

Zhang, X., Zhang, Z., Peng, H., Wang, Z., Li, H., Duan, Y., et al. (2024). GPCR-like Protein ZmCOLD1 Regulate Plant Height in an ABA Manner. Int. J. Mol. Sci. 25, 11755. doi: 10.3390/ijms252111755

Zhang, Z., and Gerstein, M. (2004). Large-scale analysis of pseudogenes in the human genome. Curr. Opin. Genet. Dev. 14, 328–335. doi: 10.1016/j.gde.2004.06.003

Zhang, Z., Ma, X., Liu, Y., Yang, L., Shi, X., Wang, H., et al. (2022). Origin and evolution of green plants in the light of key evolutionary events. J. Integr. Plant Biol. 64, 516–535. doi: 10.1111/jipb.13224

