## Supplementary material for "Evolutionary and Structural Bioinformatics Reveal GPR89 as a Conserved Solute Carrier Transporter": SUPPLEMENTARY MATERIAL.docx

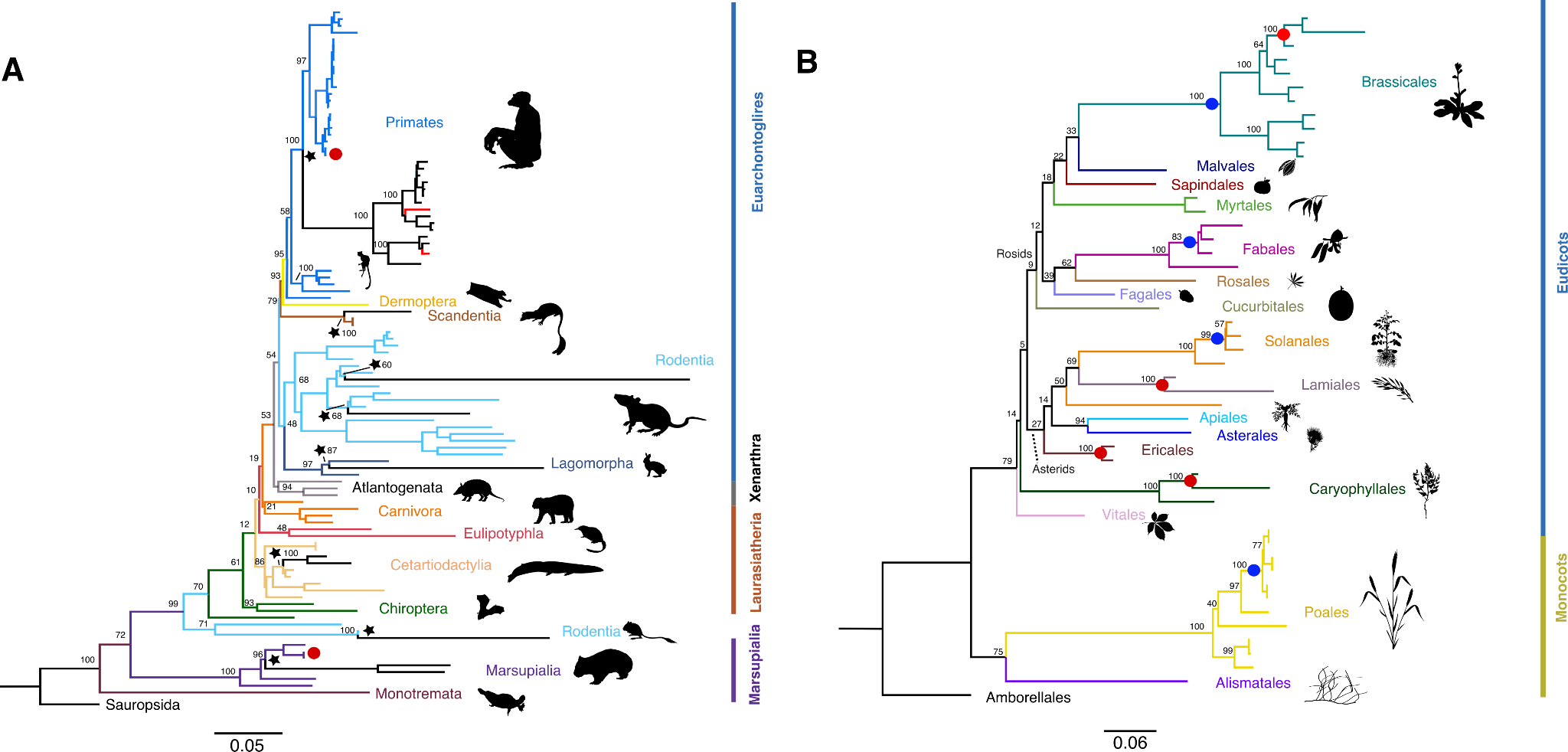

**Supplementary Figure 1: Best Maximum likelihood phylogeny representative of orthologs and paralogs of the GPR89 genes on Mammals and Angioespermae**. **(A)** Red dots indicate paralogs of GPR89 while black stars indicate GPR89P. Clack branches contain GPR89P among mammals and red branches display the presence of coding retrogenes among primates. The tree was rooted with Sauropsida GPR89gene as an outgroup. **(B)** Blue dots correspond to interspecific gene duplications while red dots indicate intraspecific gene duplication. The tree was rooted with Amborella trichopoda`s gene as an outgroup. Branch colors correspond to their respective taxonomic order. Vertical lines correspond to the taxonomic superorder. Numbers on the nodes represent Bootstrap support values. Bar indicates nucleotide substitution per site.

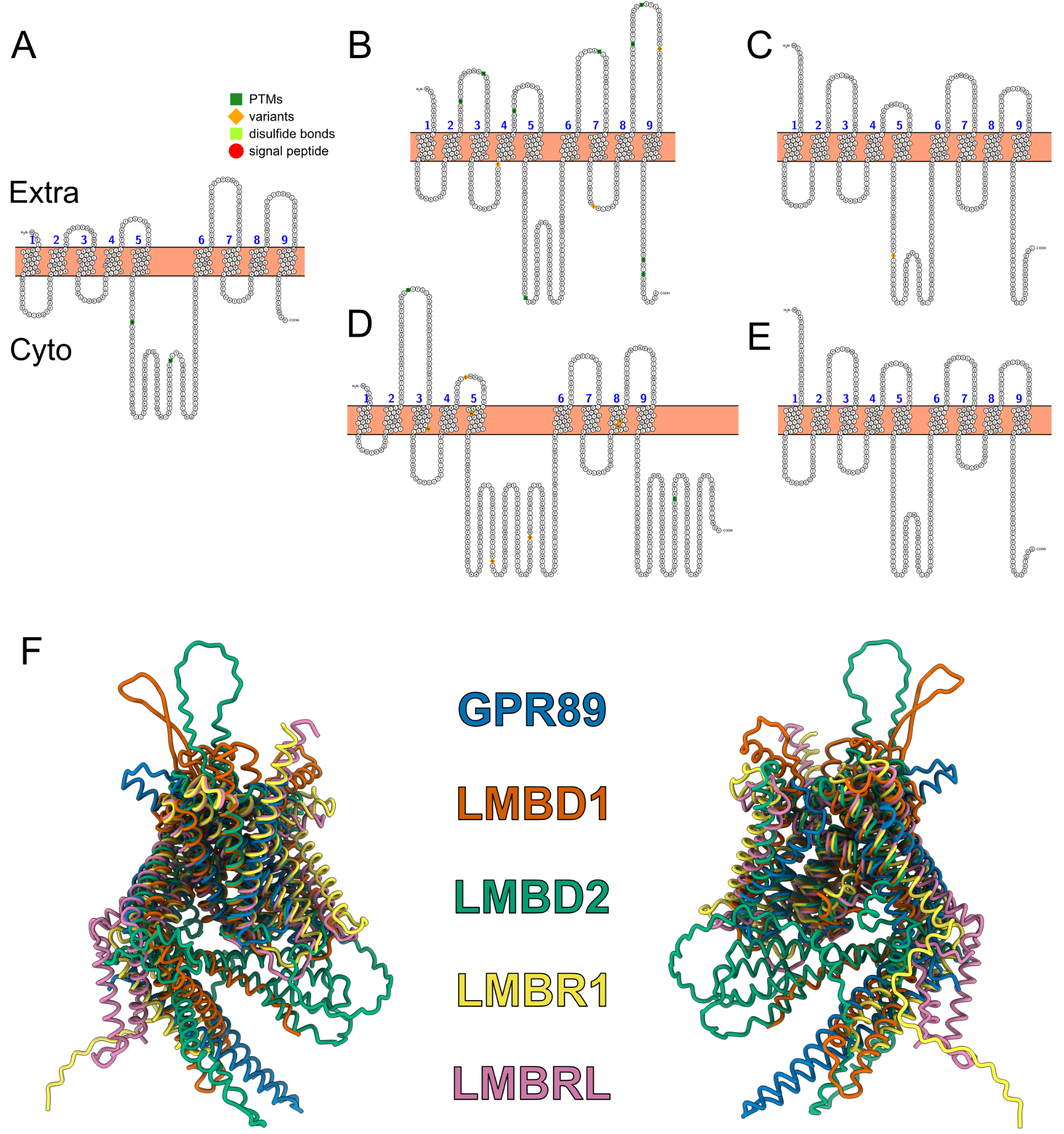

**Supplementary Figure 2:** **Topology of human LMB orthologs and GPR89.** Topological snake plot for **A)** GPR89, **B)** LMBD1, **C)** LMBR1, **D)** LMBD2 & **E)** LMBRL. Plotted using the Protter web server (Omasits *et al.,* 2014), with Uniprot accession codes. **F)** Superposed structural predictions for all human LMB orthologs & GPR89.

***Supplementary Table 1:*** **Details of the CDS sequences used on the phylogenetic analysis shown in Figure1.** Details include scientific species name, the predicted number of transmembrane segments, CDS length, number of exons, protein length and the accession number for Ensembl or NCBI databases in each case.

***Supplementary Table 2:*** **Details of the CDS sequences used on the phylogenetic analysis shown in Supplementary Figure 1.** Information about scientific species name, predicted number of transmembrane segments, CDS length, number of exons, protein length, accession number for Ensembl or NCBI databases and chromosome location for the CDS used in our second phylogenetic analysis. For the case of pseudogenes we did not report TM prediction or protein length.
